# Gut Microbiome Signatures During Acute Infection Predict Long COVID

**DOI:** 10.1101/2024.12.10.626852

**Authors:** Isin Y. Comba, Ruben A. T. Mars, Lu Yang, Mitchell Dumais, Jun Chen, Trena M. Van Gorp, Jonathan J. Harrington, Jason P. Sinnwell, Stephen Johnson, LaRinda A. Holland, Adam K. Khan, Efrem S. Lim, Christopher Aakre, Arjun P. Athreya, Georg K. Gerber, Jack C. O’Horo, Konstantinos N. Lazaridis, Purna C. Kashyap

## Abstract

Long COVID (LC), manifests in 10-30% of non-hospitalized individuals post-SARS-CoV-2 infection leading to significant morbidity. The predictive role of gut microbiome composition during acute infection in the development of LC is not well understood, partly due to the heterogeneous nature of disease. We conducted a longitudinal study of 799 outpatients tested for SARS-CoV-2 (380 positive, 419 negative) and found that individuals who later developed LC harbored distinct gut microbiome compositions during acute infection, compared with both SARS-CoV-2–positive individuals who did not develop LC and negative controls with similar symptomatology. However, the temporal changes in gut microbiome composition between the infectious (0–1 month) and post-infectious (1–2 months) phases was not different between study groups. Using machine learning, we showed that microbiome composition alone more accurately predicted LC than clinical variables. Including clinical data only marginally enhanced this prediction, suggesting that microbiome profiles during acute infection may reflect underlying health status and immune responses thus, help predicting individuals at risk for LC. Finally, we identified four LC symptom clusters, with gastrointestinal and fatigue-only groups most strongly linked to gut microbiome alterations.

## Introduction

Long COVID (LC) affects approximately 10–30% of non-hospitalized individuals infected with SARS-CoV-2, with nearly 18% of U.S. adults reporting previous history^1,2^. This condition inflicts significant morbidity, workforce loss, and an estimated economic impact of $3.7 trillion in the U.S.^3^. LC manifests through a broad spectrum of persistent symptoms and host physiological changes, ranging from cardiovascular and gastrointestinal disturbances to cognitive impairments and neurological signs and symptoms.

Notably, these symptoms are similar to those of myalgic encephalomyelitis/chronic fatigue syndrome and other post-infectious syndromes, underscoring their common clinical presentation^2,4,5^. The pathobiology of LC is complex and not well understood, complicated by the disease’s heterogeneity and emerging evidence of distinct clinical subphenotypes^6–8^.

Proposed underlying mechanisms for LC include reduced peripheral serotonin levels^9^, exaggerated humoral immune responses^10^, coagulation abnormalities^11^, neuroinflammation, persistence of viral antigens^2^, and the presence of SARS-CoV-2 in reservoir tissues^12^. The gut microbiome’s established role in shaping both innate and adaptive immunity^13^, its influence on antiviral immunity via inflammasome pathways in respiratory infections such as influenza A^14^, and may thus be of importance in LC pathogenesis.

Prior studies found an association between LC and the gut microbiome composition both during acute infection and up to six months after the infection^15–20^. However, these LC microbiome studies have focused primarily on hospitalized patients with moderate to severe illness, for whom hospitalization itself can cause shifts in the gut microbiome through altered host status^21^. Further, the lack of comparable control groups with similar symptomatology and availability of key clinical metadata such as antibiotic use, baseline comorbidities in current studies presents substantial challenges in controlling for confounders that affect microbiome composition and function, thus limiting the generalizability of study outcomes^22^. Moreover, as patient demographics shift from severely ill hospitalized individuals to outpatient populations, studying those with milder illnesses becomes more relevant, as they likely represent a broader spectrum of those at risk for LC than what has been captured in previous studies.

To address this gap, we designed a study that includes outpatients diagnosed with COVID as well as contemporaneous controls who tested negative for COVID. We found gut microbiome at the time of acute infection is a strong predictor for the development of LC and distinct microbes are associated with symptom-based LC sub phenotypes. The ability to identify patients that are likely to get LC is crucial for advancing and improved understanding of its pathobiology, the development of targeted diagnostic tools, and microbiome-based therapies to alleviate chronic symptoms.

## Results

### Study population baseline characteristics

Our study analysis comprised 947 stool samples from 799 subjects — 380 testing positive and 419 testing negative for SARS-CoV-2. Within the one-year follow-up period, 80 patients who were positive for SARS-CoV-2 developed LC, hence for downstream analysis we categorized SARS-CoV-2-positive patients as LC (SARS-CoV-2-positive patients who subsequently developed LC) or non-LC (SARS-CoV-2-positive patients who did not develop LC) A description of the study cohort census is provided in the **Methods** and **Fig. 1**. Baseline characteristics of SARS-CoV-2-positive (both LC and non-LC) and -negative participants are outlined in **Tables 1,** and **S1**. Post hoc analysis showed a higher proportion of females in the LC group than in other groups (pairwise Fisher’s exact test, *p*<0.05) and a higher number of baseline comorbidities (Dunn’s test, Bonferroni correction, *p*=0.009) compared with non-LC. The SARS-CoV-2-negative group was significantly older (Tukey’s HSD, *p*<0.001) and had a higher rate of 30-day antibiotic use (17% vs. 6%, pairwise Fisher’s exact test, *p*<0.001) compared with non-LC and higher vaccination rates (pairwise Fisher’s exact test, *p*<0.001) than other groups. Subsequent analyses were adjusted for these variables. No significant differences in BMI or race/ethnicity were observed across groups. In the SARS-CoV-2-negative group, 56% of patients were symptomatic, with sore throat (32%), cough (31%), and headache (31%) as the most common symptoms. On the other hand, 89% of the non-LC group and 99% of the LC group were symptomatic, presenting predominantly with cough (**Table S2**).

**Figure 1.**
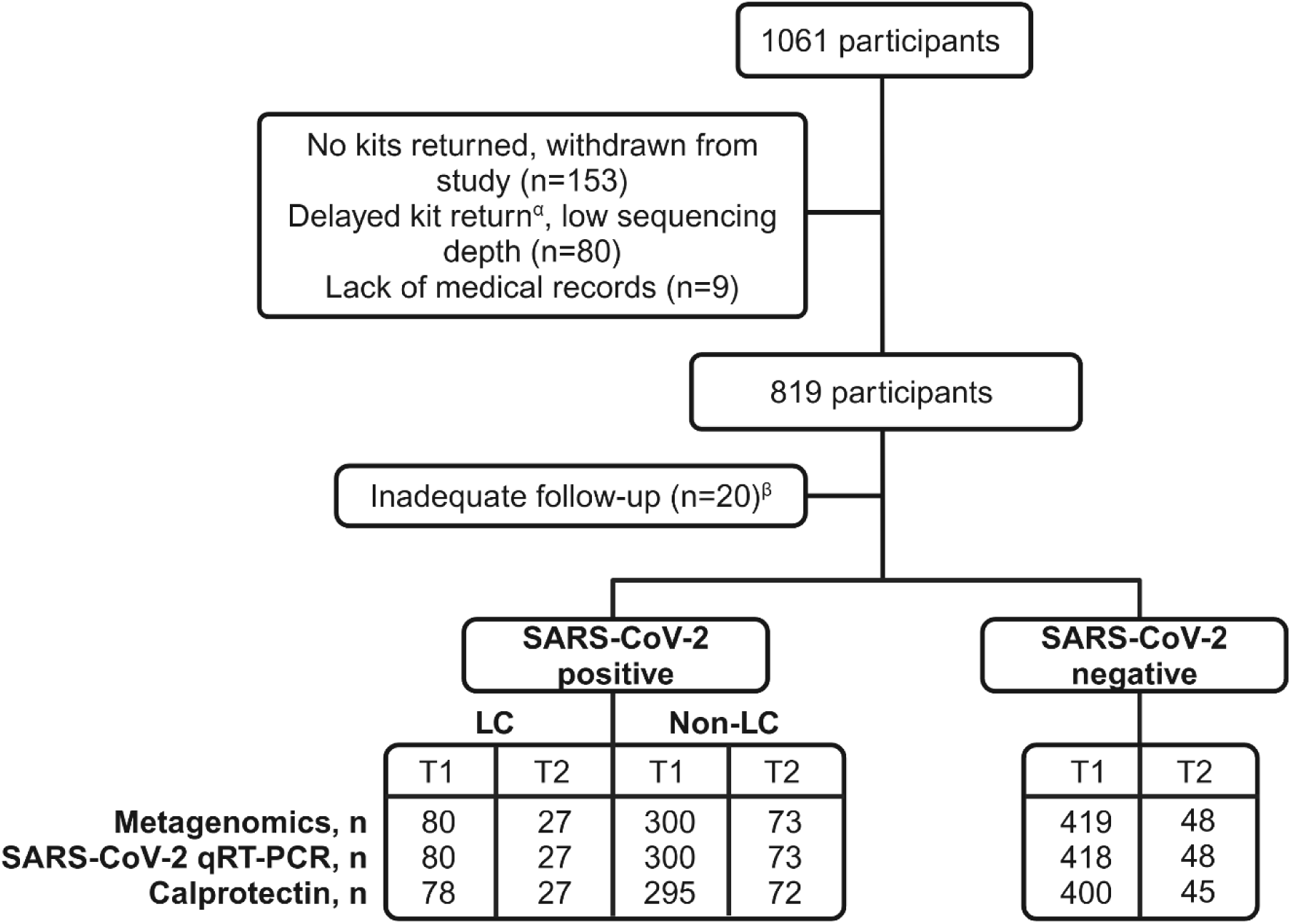
Overview of study design and number of fecal samples analyzed. Our study included three groups: Long COVID (LC), SARS-CoV-2-positive but non-long COVID (non-LC), and SARS-CoV-2-negative. Fecal samples were collected at two timepoints after the initial SARS-CoV-2 test date, denoted as T1 and T2, and were analyzed as indicated. α, The first kit was received 31 days after the initial SARS-CoV-2 test date or only second kit is returned. β, Patients in the SARS-CoV-2 positive arm with inadequate follow-up to ascertain LC status were excluded from further analysis.

**Table 1.** Baseline characteristics of study population.

|  | LC (n=80) | Non-LC (n=300) | SARS-CoV-2 negative (n=419) | *p* |
| --- | --- | --- | --- | --- |
| Age, y, mean (SD) | 56 (13) | 56 (15) | 60 (13) | <0.001 |
| Female, n (%) | 61 (76) | 169 (56) | 254 (61) | 0.005 |
| BMI, kg/m<sup>2</sup>, mean (SD) | 29.1 (6.4) | 28.6 (5.5) | 28.2 (6.4) | 0.4 |
| Race/ethnicity, n (%) |  |  |  |  |
| Hispanic (all races) | 4 (5) | 9 (3) | 15 (4) |  |
| White, non-Hispanic | 69 (86) | 283 (94) | 376 (90) |  |
| Black, non-Hispanic | 3 (4) | 2 (1) | 10 (2) |  |
| Asian, non-Hispanic | 2 (2.5) | 3 (1) | 15 (4) |  |
| All other, non-Hispanic | 2 (2.5) | 2 (1) | 2 (0.5) |  |
| Choose not to disclose/missing | - | 1 (0.3) | 1 (0.2) |  |
| CCI score, median (IQR) | 1 (0-2) | 0 (0-1) | 0 (0-2) | 0.005 |
| Symptomatic, n (%) | 79 (99) | 268 (89) | 234 (56) | <0.001 |
| Thirty-day antibiotic use, n (%) | 9 (11) | 17 (6) | 70 (17) | <0.001 |
| Fully vaccinated, n (%) | 18 (23) | 95 (32) | 298 (71) | <0.001 |

Abbreviations: BMI, body mass index; CCI, Charlson Comorbidity Index; IQR, interquartile range; SD, standard deviation. CCI scores and time to kit receipt were analyzed using the Kruskal–Wallis test, with post hoc assessment via Dunn's test with Bonferroni correction. Age and BMI were evaluated with an unbalanced one-way ANOVA, and age differences were further analyzed using Tukey's HSD. Sex, 30-day antibiotic use, and vaccination status were assessed with Fisher's exact test, followed by pairwise Fisher's exact test with Holm correction for post hoc analysis. In the SARS-CoV-2-negative group, BMI for five patients, antibiotic use, and CCI scores for three patients were missing.

### Long COVID patients harbor a distinct gut microbiome during acute infection

Alpha diversity based on the Chao1 metric is significantly lower in SARS-CoV-2 positive patients (both LC and non-LC) compared to SARS-CoV-2-negative subjects but comparable based on Shannon index (**Fig. 2A**). PCoA based on Bray–Curtis (BC; PERMANOVA, R²=0.004, *p*<0.001; **Fig. 2B**) and GUniFrac (PERMANOVA, R²=0.004, *p*=0.002; **Fig. S1**) beta diversity metrics show significant differences in microbial composition among SARS-CoV-2-positive patients who develop LC, those who do not develop LC, and SARS-CoV-2-negative subjects, after adjusting for age, sex, baseline comorbidities, and antibiotic use. Post hoc analysis reveals that patients who develop LC have a distinct gut microbiome profile during acute infection compared to those who do not develop LC and SARS-CoV-2-negative subjects (pairwise comparisons using PERMANOVA, FDR-adjusted *p*<0.05). Furthermore, during acute infection, the microbial diversity of patients who develop LC or do not develop LC falls along two RDA axes, with changes of similar magnitude occurring in different directions, indicating distinct microbial diversity gradients (**Fig. 2C**). Patients who develop LC exhibit significantly higher levels of *Faecalimonas sp000209385, Anaerovoracaceae; CAG-145 sp900545135, Blautia sp900539145, Schaedlerella glycyrrhizinilytica, Massilimaliae timonensis, Ruthenibacterium lactatiformans, Faecalimonas sp900550235, Acutalibacteraceae; UBA945 sp900755905, and Eubacterium G sp900552275,* while those who do not develop LC have significantly higher levels of *Faecalibacillus intestinalis, Amedibacillus dolichus, Blautia A obeum, Blautia A sp900548245, Slackia A sp900553655, Faecalibacillus faecis, Slackia A sp900553655, Faecalibacillus sp900544435, Acutalibacteraceae; CAG-964 sp000435335, Collinsella sp900551635, Blautia A sp900120195, Blautia A sp900551715, Collinsella sp900554455, Blautia A sp900549015, Blautia A obeum B, Eubacterium G sp900552275, Anaerostipes sp900066705, lntestinibacter sp900553485,* and *Anaerobutyricum sp900554965* compared to SARS-CoV-2-negative subjects at the time of infection with minimal to no overlap between taxa (ZicoSeq, FDR-adj. *p*<0.1; **Fig. 2D**).

**Figure 2.**
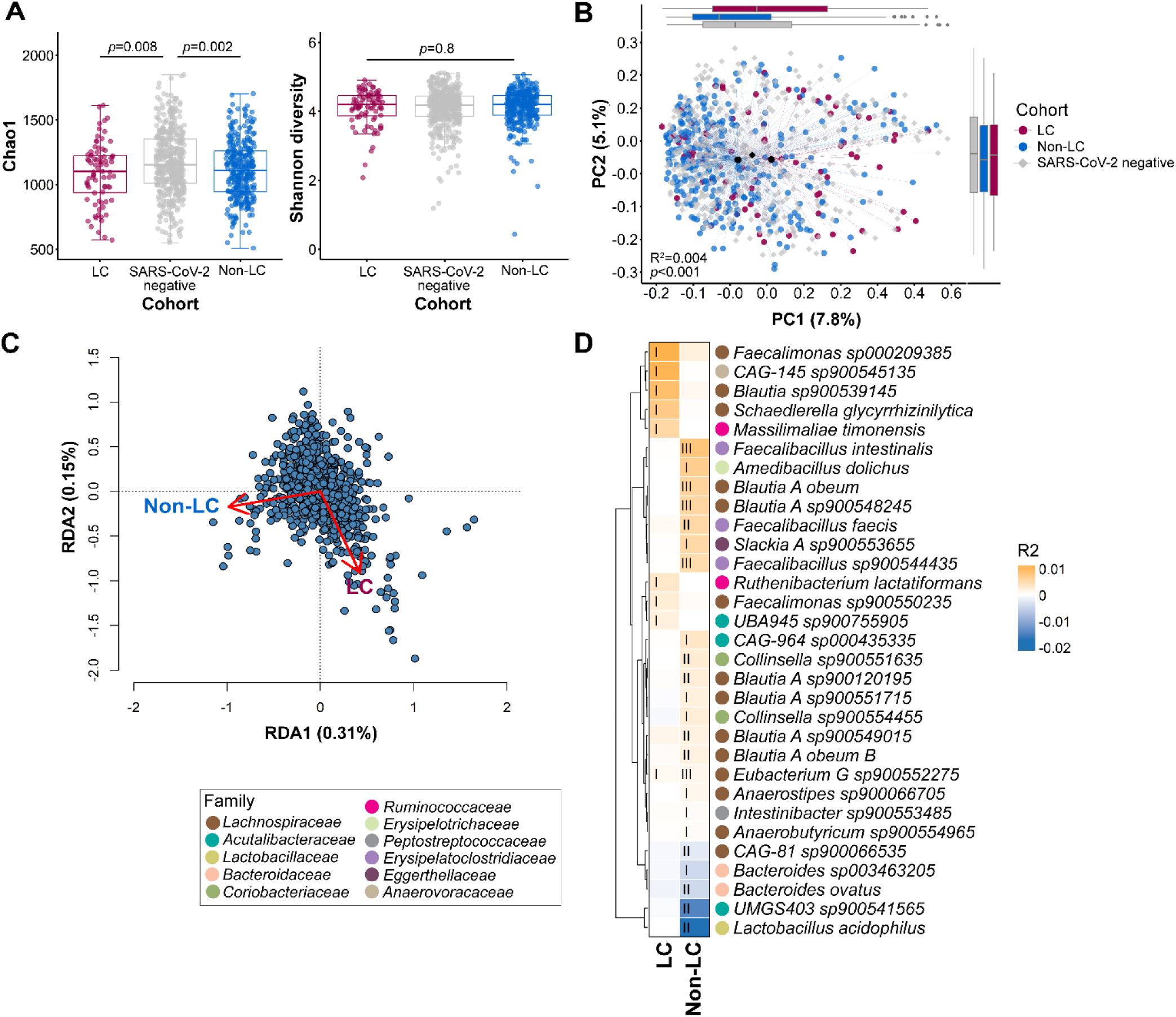
LC patients harbor a distinct baseline gut microbiome compared with non-LC and SARS-CoV-2-negative individuals. **(A)** Alpha diversity plotted using the Chao1 and Shannon indices. **(B)** Principal coordinate analysis (PCoA) of species-level Bray–Curtis distances of LC (n=80) vs. non-LC (n=300) subjects and SARS-CoV-2-negative controls (n=419) adjusted for age, sex, baseline Charlson Comorbidity Index (CCI) score, and antibiotic use. **(C)** Redundancy analysis (RDA) reveals the directionality and effect size of LC and non-LC group memberships on species-level microbial diversity of samples collected during acute infection. **(D)** Heatmap displaying differential species-level abundances between SARS-CoV-2-negative controls (n=419) and LC patients (n=80, adjusted for age and sex) or non-LC patients (n=300, adjusted for age, CCI score, and antibiotic use). Levels of statistical significance are represented by Roman numerals: I, FDR-adjusted *p*<0.1; II, *p*<0.05; and III, for *p*<0.01.

Abundances of microbial genes assorted into pathways significantly differed among the three groups (BC dissimilarities, PERMANOVA, R^2^=0.009, *p*=0.002), but these differences diminished after adjusting for clinical confounders (R^2^=0.004, *p*=0.09). Compared to SARS-CoV-2 negative controls, the non-LC cohort exhibited significant reductions in 18 microbial pathways and enrichments in 3 pathways (ZicoSeq, FDR-adjusted *p*<0.1; **Fig. S3A**). Specifically, pathways related to nitrogen cycling and amino acid metabolism—including the urea cycle, anaerobic purine nucleobase degradation II, L-histidine degradation III, and L-citrulline biosynthesis—were significantly decreased in non-LC individuals. In contrast, pathways involved in carbohydrate degradation and cofactor biosynthesis demonstrated variable trends between non-LC and SARS-CoV-2 negative cohorts while no statistically significant differences in mapped and integrated microbial pathways were observed between LC and SARS-CoV-2 negatives **(Fig. S3A, B**).

Fecal calprotectin, a well-established biomarker for gut inflammation and neutrophil activity, is a potential indicator of these signs in COVID-19 patients^23,24^. However, in our cohort, fecal calprotectin levels were not associated with SARS-CoV-2 positivity or LC at any study phase (Fisher’s exact test, *p≥*0.7 for both time points; **Table 2**). During acute infection, the PCR detection rate of the SARS-CoV-2 E gene in stool samples was 11% for the SARS-CoV-2-positive cohort (LC and non-LC), with 7.5% and 12% positivity rates in the LC and non-LC groups, respectively. We also saw no association between fecal PCR positivity and calprotectin level (Fisher’s test, *p*=0.55). Among the successfully sequenced SARS-CoV-2-positive samples, B.1.2 (n=11) was the most prevalent lineage, followed by B.1.617.2 (delta variant, n=5), and AY.103 (n=3) (**Table S3**).

**Table 2.**
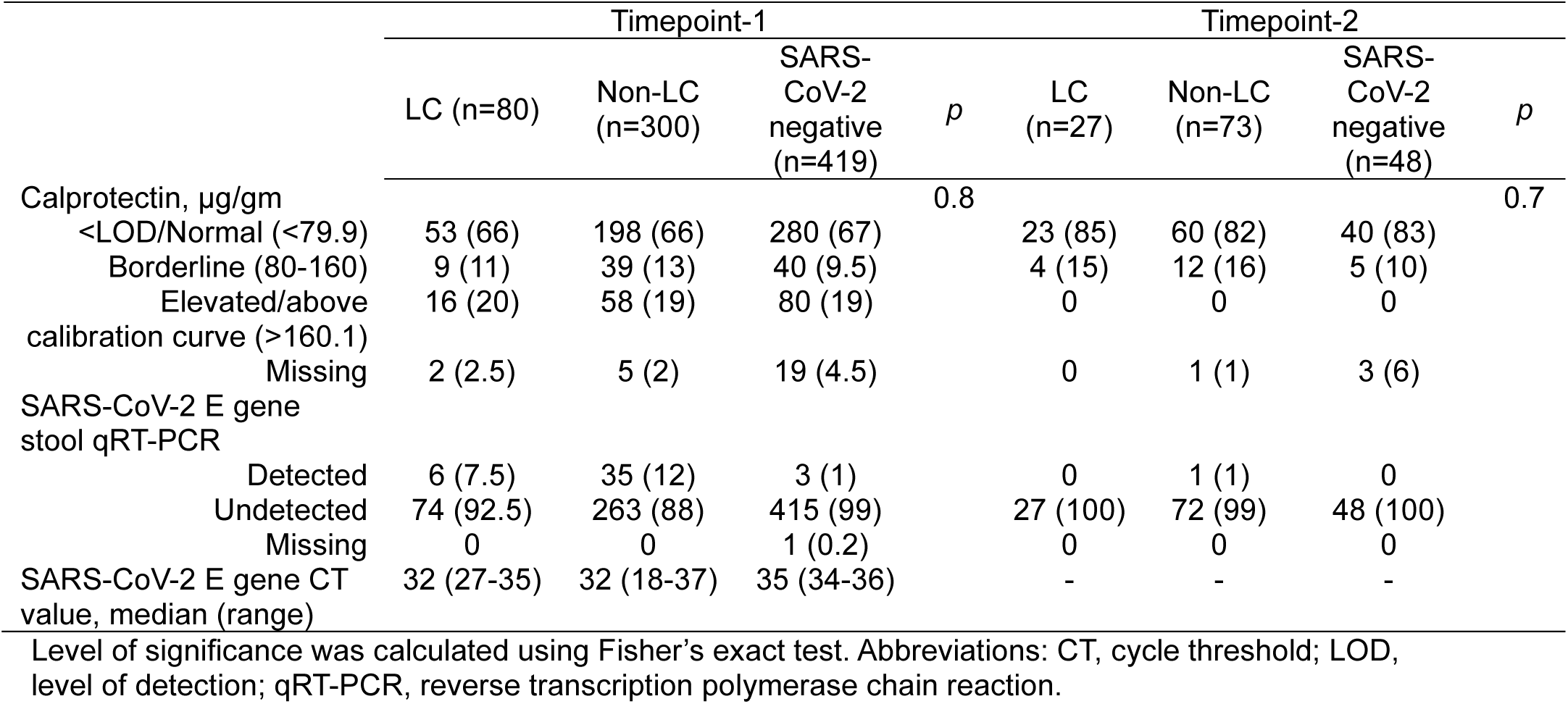
Qualitative stool calprotectin levels and SARS-CoV-2 qRT-PCR characteristics.

| | Timepoint-1 | | | $p$ | Timepoint-2 | | | $p$ |
| --- | --- | --- | --- | --- | --- | --- | --- | --- |
|  | LC (n=80) | Non-LC (n=300) | SARS-CoV-2 negative (n=419) |  | LC (n=27) | Non-LC (n=73) | SARS-CoV-2 negative (n=48) |  |
| Calprotectin, $\mu\text{g/gm}$ | | | | 0.8 | | | | 0.7 |
| <LOD/Normal (<79.9) | 53 (66) | 198 (66) | 280 (67) |  | 23 (85) | 60 (82) | 40 (83) |  |
| Borderline (80-160) | 9 (11) | 39 (13) | 40 (9.5) |  | 4 (15) | 12 (16) | 5 (10) |  |
| Elevated/above calibration curve (>160.1) | 16 (20) | 58 (19) | 80 (19) |  | 0 | 0 | 0 |  |
| Missing | 2 (2.5) | 5 (2) | 19 (4.5) |  | 0 | 1 (1) | 3 (6) |  |
| SARS-CoV-2 E gene stool qRT-PCR |  |  |  |  |  |  |  |  |
| Detected | 6 (7.5) | 35 (12) | 3 (1) |  | 0 | 1 (1) | 0 |  |
| Undetected | 74 (92.5) | 263 (88) | 415 (99) |  | 27 (100) | 72 (99) | 48 (100) |  |
| Missing | 0 | 0 | 1 (0.2) |  | 0 | 0 | 0 |  |
| SARS-CoV-2 E gene CT value, median (range) | 32 (27-35) | 32 (18-37) | 35 (34-36) |  | - | - | - |  |

Level of significance was calculated using Fisher's exact test. Abbreviations: CT, cycle threshold; LOD, level of detection; qRT-PCR, reverse transcription polymerase chain reaction.

### Gut microbiome composition does not change significantly over time at the cohort level, independent of LC status

We then analyzed the temporal variability in gut microbiome composition over the two time-points of sample collection after SARS-CoV-2 testing. Sampling timelines and median return times for kits are outlined in **Fig. 3A**. Time to kit receipt did not differ between groups for both timepoints (Tukey HSD, *p*>0.1). Principal coordinate analysis (PCoA) of BC dissimilarities showed significant changes in microbial composition over time within individuals (PERMANOVA, within-individual permutation, *p*=0.001; **Fig. 3B**) and within each cohort (**Fig. 3C**). However, the extent of these changes does not significantly differ among the three cohorts (Tukey’s HSD *p*>0.5 for all comparisons; **Fig. 3D**) suggesting that the changes among the two time points represents the normal variability expected over time and is unlikely to underlie the development of LC.

**Figure 3.**
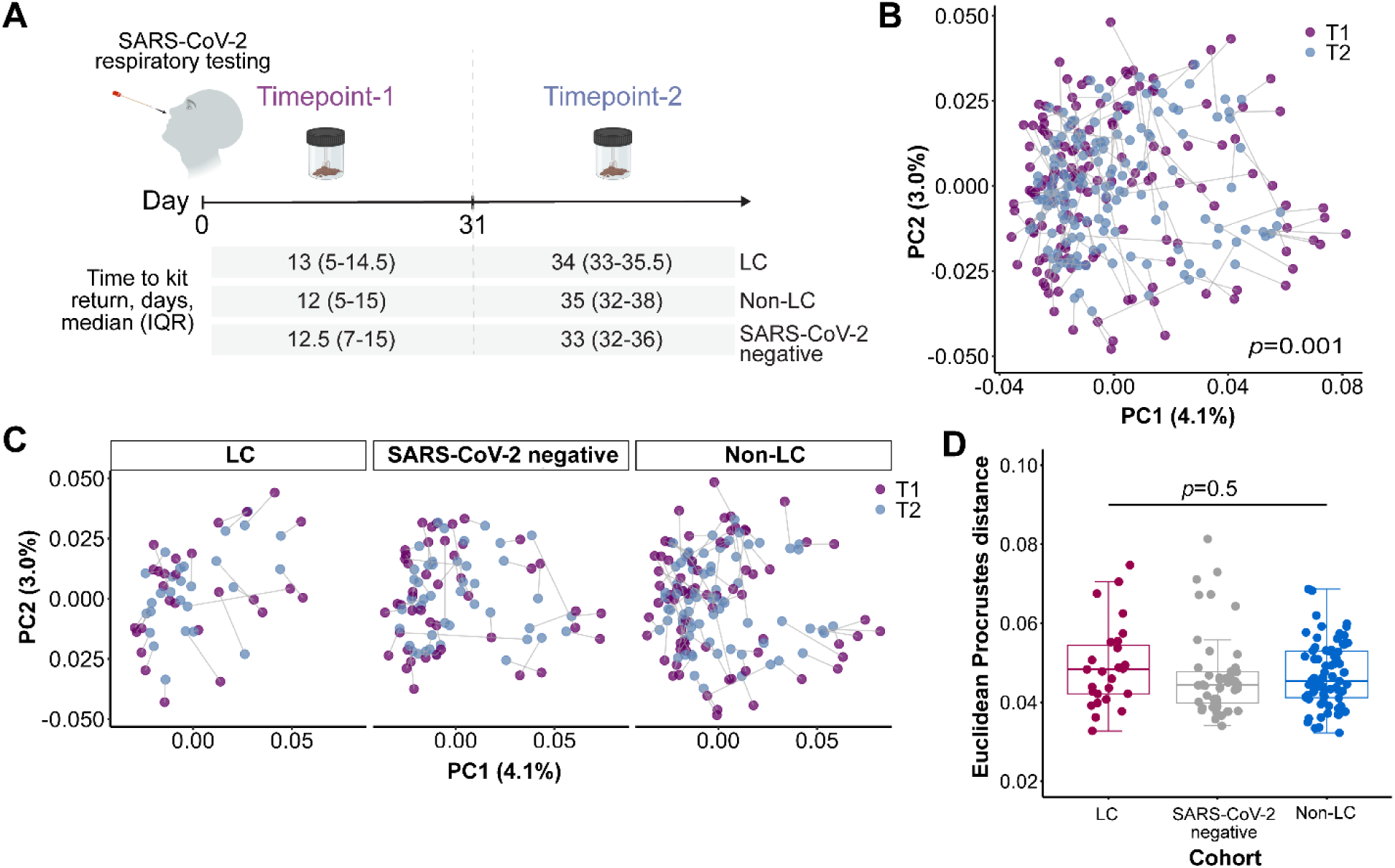
Longitudinal gut microbiome variability across study cohorts. **(A)** Overview of stool sampling timelines and median return times for kits. PCoA plots using Bray–Curtis dissimilarities were generated to compare microbial communities within individual subjects **(B)** and across different cohorts **(C)** between the two sampling timepoints. Timepoint 1 and 2 samples from the same subject are interconnected by lines. Procrustes analysis showed statistically significant similarity in the spatial arrangement of microbial communities between timepoint 1 and 2 samples (*p*=0.001). **(D)** Boxplots illustrating Euclidean distances calculated from pairwise Procrustes distances across three study cohorts. No statistically significant differences were noted (Tukey’s HSD *p*≥0.5 for all comparisons). Numbers of patients: LC (n=27), non-LC (n=73), and SARS-CoV-2-negative (n=48).

### Gut microbiome composition during acute infection predicts long COVID

To determine the ability of gut microbiome to predict LC in 80 individuals from a cohort of 380 SARS-CoV-2-positive patients, we employed ML algorithms — logistic regression with L1 regularization, and ANN (**Fig. 4A**). We initially validated models that use clinical variables or species- and genus-level bacterial abundances separately. We found gut microbiome composition alone at the species level or the genus level achieved a higher accuracy than clinical variables alone (Student’s *t*-test, *p*<0.05; **Fig. 4B**). To investigate whether clinical features improve the predictive performance of the models using microbiome data, we then added clinical variables to microbiome data which resulted in marginal improvements in accuracy (up to 2%).

**Figure 4.**
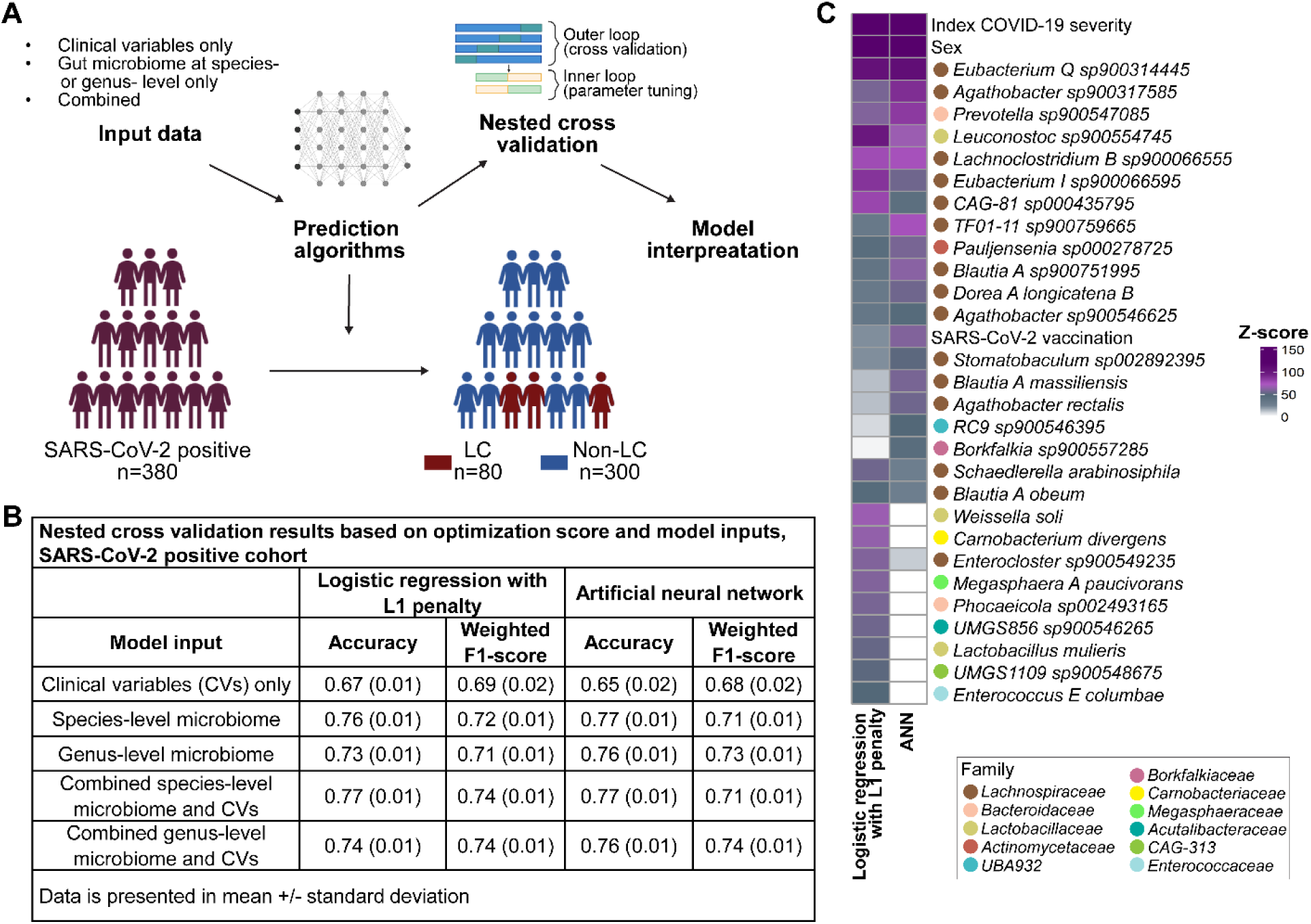
Gut microbiome composition and clinical variables can predict LC in a SARS-CoV-2 positive cohort. **(A)** Framework for developing and validating predictive algorithms — logistic regression with L1 regularization, and artificial neural network (ANN) for predicting LC within a cohort of COVID-19 patients. Model inputs included clinical variables, and species- and genus-level microbiome abundances, as well as their integration, to calibrate each model’s predictive power. **(B)** Nested cross-validation results per model inputs and model types. **(C)** Heatmap showing frequencies of the top 20 most impactful features over 50 nested cross-validation iterations, optimized for accuracy. This curated list of features consistently influenced model outcomes. Additional information is provided in the **Supplemental Methods**.

To evaluate the consistency of model decisions across 50 nested cross-validation runs, we identified the 20 predictors with the highest demonstrable predictive performance and defined as the most impactful features for each run. Overall, from 5,679 input features used in the model across the cross-validation runs, we identified 107 features in logistic regression, and 175 in ANN, as the most impactful features. Across each model run, the most impactful clinical predictors for LC were index COVID-19 disease severity, and sex. In ANN, these were followed by *Eubacterium Q sp900314445, Agathobacter sp900317585, Prevotella sp900547085, Lachnoclostridium B sp900066555, Lachnospiraceae; TF01-11 sp900759665, Leuconostoc sp900554745, and Blautia A sp900751995*. In logistic regression with L1 regularization, most impactful microbial predictors included mainly *Eubacterium Q sp900314445, Leuconostoc sp900554745, Lachnospiraceae; CAG-81 sp000435795, Eubacterium I sp900066595*, *Lachnoclostridium B sp900066555, Weissella soli, and Carnobacterium divergens* (**Fig. 4C**). Among the most impactful features across models, 68 predictors were shared between ANN (39%) and logistic regression (63.5%), 63 of which were microbial predictors, including 31 species from *Lachnospiraceae* family (**Supplemental data)**. The remaining five were clinical covariates including index COVID-19 severity, sex, SARS-CoV-2 vaccination status, immunosuppression and thirty-day antibiotic use.

### Symptom based Long COVID subphenotypes are associated with distinct gut microbial composition

Long COVID is a heterogeneous entity encompassing symptoms across multiple organ systems. We found fatigue was the most prevalent symptom, reported by 51 individuals, followed by dyspnea (n=36) and cough (n=27). These symptoms were not mutually exclusive. For example, among the 80 patients with LC, 26 (32.5%) experienced only one symptom, while 13 (16%) reported two or three symptoms, and 41 (51%) had more than three symptoms. A subset of patients (16%, n=13) required frequent clinical visits and received specialized care from clinics dedicated to LC management. While gut microbiome has been associated with the development of LC, it remains unclear if there are specific symptoms or groups of symptoms that are more likely to be driven by changes in gut microbiome. We identified four distinct LC subphenotypes based on symptom co-occurrence. Interestingly these aligned with different organ systems (**Fig. 5A**) and include (1) Gastrointestinal & sensory: Abdominal pain, diarrhea, anosmia/dysgeusia. (2) Musculoskeletal & neuropsychiatric: Myalgia, arthralgia, impaired mobility, paresthesia, headache, lightheadedness, brain fog, sleep problems, palpitation, and mood issues. (3) Cardiopulmonary: Dyspnea, chest pain, exertional malaise, and cough. (4) Fatigue-only: Fatigue.

**Figure 5.**
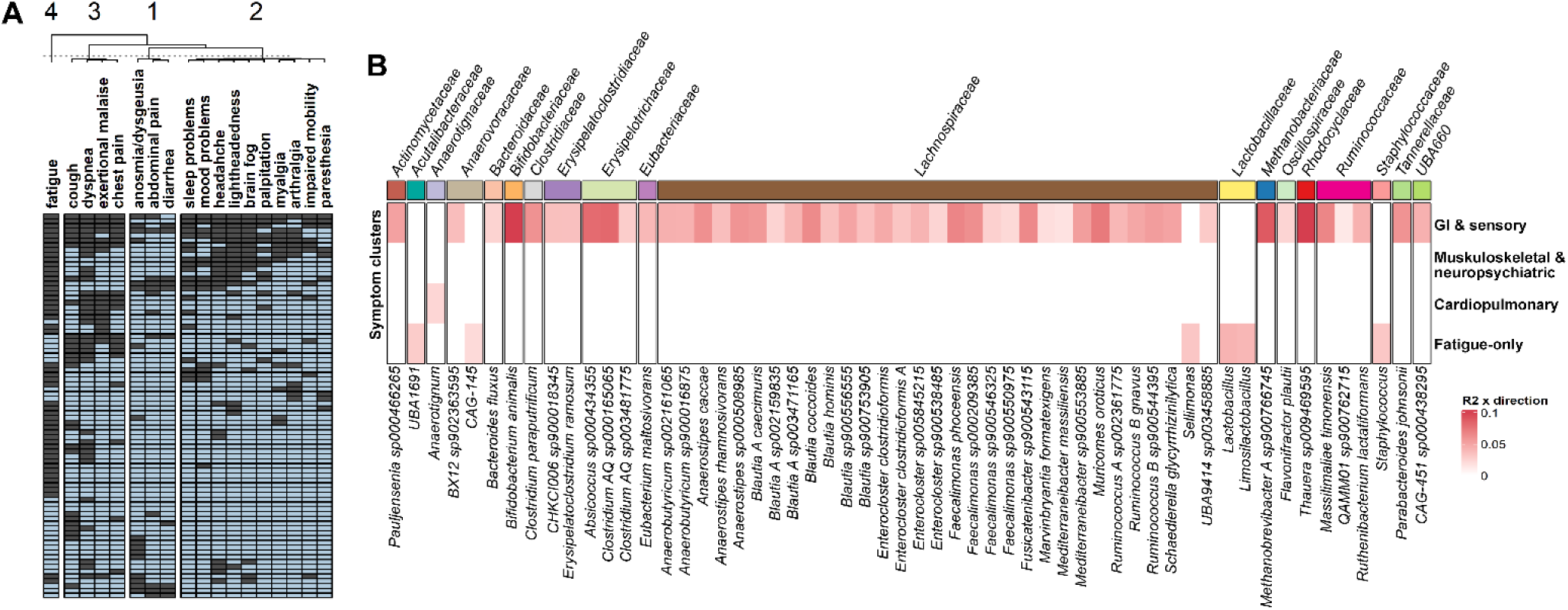
Data-driven clustering of LC symptoms reveals distinct disease subphenotypes associated with unique microbial signatures. **(A)** Hierarchical and *k*-means clustering of LC symptoms observed in at least 10% of patients revealed four distinct symptom clusters. **(B)** Differential taxa abundances at the species and genus levels were analyzed between SARS-CoV-2 positive patients with and without individual symptom clusters. Multivariate analysis using ZicoSeq (square-root transformation), adjusting for sex in the LC group and for the remaining symptom clusters in each cluster analysis. Taxa with FDR-adjusted *p*<0.1 are shown.

To determine the potential role of gut microbiome in the different LC subphenotypes, we first defined symptom cluster memberships for LC patients based on the presence of one or more cluster-specific symptoms. Following this classification, we employed ZicoSeq to analyze species-level gut microbiome composition associated with these symptom clusters. In this analysis, we adjusted for untested clusters and sex. We found that the gastrointestinal and sensory cluster exhibited the most prominent changes in taxa abundance with 50 species, including 30 from *Lachnospiraceae* family, 3 each from *Anerovoracaeae*, *Erysipelatoclostridiaceae*, *Erysipelotrichaceae*, and *Ruminococcaceae* families significantly increased in comparison with SARS-CoV-2 positive patients who do not develop persistent GI/sensory symptoms (FDR adj. *p*<0.1; **Fig. 5B**). Interestingly, 20 of these 50 species were also significantly increased when the LC group was compared to the non-LC cohort, suggesting that the overall compositional differences could be driven by this symptom cluster (**Supplementary data**). Specifically, fatigue-only group was associated with multiple genera including *Lactobacillus*, *Limosilactobacillus*, *Sellimonas*, *Staphylococcus*, *Acutalibacteraceae; UBA1691* and *Anaerovoracaceae; CAG-145*, whereas there were minimal associations with cardiopulmonary, and musculoskeletal & neuropsychiatric clusters.

## Discussion

In this study, we found that SARS-CoV-2-positive individuals who later developed LC harbored a distinct gut microbiome composition at the time of acute infection. Although prior studies provide insight into alterations in gut microbiome and their associations with COVID-19 outcomes, they offer few insights into the ability of the gut microbiome at the time of acute infection to predict subsequent LC especially in outpatient cohorts. To address this knowledge gap, we leverage our unique longitudinal design, to assess the predictive power of the gut microbiome and/or clinical features using various ML models, including artificial neural networks (ANN) and logistic regression with L1 regularization. Our analyses revealed that species-level microbiome data during acute infection served as a stronger predictor of long COVID development compared to clinical variables alone. Notably, the addition of clinical variables such as disease severity, sex, and vaccination status only marginally improved the performance of the microbiome-based model, suggesting that gut microbiome composition alone may more accurately reflect underlying health conditions and immune status.

Specific bacterial species, including *Prevotella* spp., *Leuconostoc spp.*, and members of the *Lachnospiraceae* family like *Eubacterium spp.* and *Agathobacter* spp., were identified as key contributors to the models’ predictive capabilities. Together, our findings not only corroborate previous studies that have reported correlations between the gut microbiome and long COVID but also, to our knowledge, provide the first evidence demonstrating the predictive potential of the gut microbiome during the initial infection phase. This insight could be harnessed as a diagnostic tool to identify patients at higher risk for developing long COVID, enabling early interventions and personalized treatment strategies.

There is growing recognition that long COVID is a heterogeneous disorder characterized by a wide range of symptoms, likely representing different subphenotypes. A previous study^7^ used topic modeling on newly incident conditions (30-180 days) extracted from EHR and identified four LC subphenotypes including (1) cardiac and renal, (2) respiratory, sleep and anxiety, (3) musculoskeletal and nervous and (4) digestive and respiratory. Another study conducted in China with two cross sectional and one longitudinal cohort studied the role of gut microbiome in long COVID phenotypic heterogeneity and they have found three microbiome-based enterotypes associated with disease heterogeneity^20^. Our approach differed in that we based our subphenotype identification on the co-occurrence of symptoms, leading to the identification of distinct system based long COVID symptom clusters. We then studied the microbiome composition in relation to these clusters and found that they are differentially associated with the gut microbiome. Interestingly, patients developing gastrointestinal/sensory symptoms, and fatigue as part of LC were associated with significant alterations in microbial composition at the time of acute infection while those developing cardiopulmonary and musculoskeletal & neuropsychiatric clusters had minimal associations to specific genera or species. Similar to the recent study^20^, we found changes in *Lachnospiraceae* were associated with the development of GI symptoms. Interestingly, members of *Lachnospiraceae* family are a one of the major short chain fatty acid producers with proposed impact on the host physiology and gut immunity as well as the disease and health states^25,26^.

In this study, we incorporated extensive clinical metadata alongside stool metagenomics and biomarkers of GI function across three cohorts: SARS-CoV-2-positive subjects with or without LC, and contemporaneous SARS-CoV-2-negative controls presenting with similar symptomatologies. Several strengths of our study highlight its unique contributions. Unlike prior studies that primarily focused on hospitalized COVID-19 patients, we examined a relatively large outpatient cohort with predominantly mild disease, which is more representative of the current post-pandemic population. Most previous studies included control groups composed of asymptomatic individuals, non-disease controls from colonoscopy trials, or healthy controls recruited pre-pandemic—all with minimal comorbidities and either no or unknown antibiotic usage status. These factors, which were considered in our study models, could significantly impact the gut microbiome and thus serve as confounders in earlier studies. A key strength of our study was the inclusion of a contemporary control group, enabling robust adjustment and control of microbiome analyses for clinical confounders.

Our rigorous methodology, including ML models and subphenotyping, further strengthens our findings. In contrast to methods that rely on ICD codes to identify symptoms, which often emphasize respiratory symptoms at the expense of other manifestations like neurological or cardiovascular issues^2^, we manually reviewed all symptoms and conditions that patients developed after SARS-CoV-2 testing. Specifically, the identification of gut microbiome signatures associated with only a few symptom clusters suggests that these symptom groups are more likely driven by alterations in gut microbiome composition. Different from previous approaches, we employed techniques to identify the specific taxa the models relied on for their decisions. These taxa were consistently shared across the models, which is important for interpretability and reliability of ML models for end users.

Despite these strengths, our study has some limitations. While this represents one of the largest cohort studies involving outpatient COVID-19 patients, the subset of LC patients (n=80) was relatively modest. Nevertheless, the two models we employed showed promise by capturing signals that could reliably predict LC using only microbiome data, as indicated by cross-validation metrics. However, to increase confidence in these findings, they require external validation. The lack of a comparable outpatient dataset with microbiome sampling during acute infection restricts our ability to validate our results using previously published studies (**Table S4**). Future studies that include a larger dataset with longitudinal sampling over a longer period can help build on our findings and validate the performance of our model.

## Conclusion

Our results demonstrate that the gut microbiome during acute SARS-CoV-2 infection can potentially identify patients at risk of developing LC. Variations in the gut microbiome may be intricately linked to the pathophysiology of specific LC symptoms, including gastrointestinal/sensory and fatigue-only clusters. Future studies incorporating the microbiome are needed to refine ML-based predictive models, further validate findings from our study, and assess potential microbiome-driven mechanisms that underlie the development of symptoms in LC patients.

## Methods

### Study design and population

The study was reviewed and approved by the Mayo Clinic Institutional Review Board (# 20-005988). We recruited adults aged 18 years or older who underwent SARS-CoV-2 respiratory testing at Mayo Clinic’s multi-site health system in Minnesota, Florida, and Arizona, from October 2020 to September 2021. Potential participants were identified by reviewing daily reports from electronic health records (EHRs) filtered by SARS-CoV-2 testing and scheduling orders. After confirming eligibility of age (18+), capacity to consent (per filing of legal authorized representatives or power of attorney), females not currently pregnant (at the time of medical record reviewed and confirmed scheduled SARS-CoV-2 test, subjects were contacted via a recruitment email. An informed consent was obtained from each participant. Of the initial 1,061 participants recruited, 242 were excluded due to failure to return kits, withdrawal from the study, delayed kit returns, failed sequencing, or incomplete medical records. The remaining 819 participants were followed prospectively with review of EHR for at least 12 months. An additional 20 participants where adequate clinical follow-up was not available were excluded. The final analysis included 947 stool samples from 799 subjects (380 SARS-CoV-2 positive and 419 SARS-CoV-2 negative) (**Fig. 1**).

### Specimen collection and data generation

#### Stool sample collection

Participants were asked to submit two stool samples at week 0-2 and week 3-5 after initial SARS-CoV-2 test. Home collection kits, along with detailed collection instructions, were mailed to each participant. Upon collection, samples were returned in tubes on frozen gel packs via overnight FedEx service. Stool samples returned within 31 days of the index date were classified as Timepoint 1 (T1) samples, and those received afterward as Timepoint 2 (T2) samples. T1 represents the acute illness (0–1 month) and was used for the majority of the downstream analyses. They were stored at −80°C till they were submitted for next-generation sequencing as outlined below and in **Supplemental Methods**.

#### Microbial DNA extraction, sequencing, and taxonomic assignment

Stool aliquots were sent to the University of Minnesota Genomics Center. DNA extraction from each sample was performed using Qiagen’s DNeasy 96 PowerSoil Pro QIAcube HT Kit following the manufacturer’s instructions. Metagenomic libraries were prepared using the Nextera XT protocol. Samples were sequenced on a NovaSeq 6000 (Illumina, San Diego, CA, USA), targeting 8 million reads per sample (2×150 bp). Taxonomic profiling of reads was performed using Kraken2 with the GTDB v202 database^27^. Relative abundances were then estimated using Bracken^28^. Functional profiling of metagenomic data was performed using HUMAnN3^29^.

#### Stool calprotectin measurement and SARS-CoV-2 RNA detection

Calprotectin in stool was measured at Arizona State University using the BÜHLMANN fCAL ELISA kit (BÜHLMANN Diagnostics Corp, Amherst, NH) per the manufacturer’s instructions. For the extraction preparation, ∼30mg of stool was weighed and transferred into a calprotectin assay tube. The samples were diluted 1:50 with the kit extraction buffer based on input weight, then were vortexed 3 minutes at 3,000×g. The samples were then heat-inactivated at 65°C for 30 minutes. Following heat inactivation, the samples were centrifuged at 3,000×g for 5 minutes. The supernatant was transferred to Eppendorf tubes and stored at −80°C until further analysis. A four-parameter logistic function, calibrated with five standard samples in each assay, was used to convert sample optical density to calprotectin concentration. Concentrations were normalized to adjust for varying stool dilution volumes. Based on the clinical threshold guidelines provided calprotectin status of each sample was categorized as normal (<80 μg/g), gray zone/borderline (80–160 μg/g), or elevated (>160 μg/g).

#### Stool SARS-CoV-2 RNA detection and SARS-CoV-2 genome sequencing

200 mg of stool was added to 1.7ml of PBS and vortexed for 3 minutes at 3,000×g. The samples were centrifuged at 20,000×g for 5 and then 1ml of supernatant was transferred into a bioMérieux eMAG (Marcy-l’Étoile, France) for total nucleic acid extraction. Detection of SARS-CoV-2 was performed by TaqMan-based RT-qPCR using SuperScript III Platinum One-Step RT-qPCR Kit (Invitrogen, Carlsbad, CA) and E-Sarbeco primers and probe as previously described^30^. A 25µl total reaction of 4.5µl molecular grade water, 12.5µl of SSIII 2x-MasterMix, 1µl of each 10µm primer, 0.5µl of 10µm probe, 0.5µl of SSIII Taq, and 5µl of TNA. Cycling conditions on a Quant Studio 7 were a 50° RT step for 5 minutes, a single denature of 95° for 2 minutes, followed by 45 cycles of 95° for 3 seconds and 58° for 30 seconds.

SARS-CoV-2 genome sequencing was performed on samples that were SARS-CoV-2 RNA-positive using the COVIDSeq Test (Illumina, San Diego, CA, USA). Libraries were sequenced on the Illumina NextSeq2000 instrument using 2×10^9^ paired end reads. Sequencing reads adapter sequences were trimmed using trim-galore, aligned to the Wuhan1 reference genome (MN908947.3) using the Burrows–Wheeler aligner, BWA-MEM version 0.7.17-r1188 and primer sequences trimmed using iVAR version 1.3.1. SARS-CoV-2 lineage calling was performed using Pangolin^31^.

### Clinical data acquisition and case definitions

LC was defined as the persistence of symptoms for three months or more after onset of illness or new-onset symptoms not explained by an alternate diagnosis^32^. The index date was defined as the SARS-CoV-2 test date for both SARS-CoV-2-positive and -negative groups. To ascertain symptom status and types experienced, a prospective review of EHRs, including provider notes, SARS-CoV-2 testing orders, nurse screening forms, and patient portal communications, was conducted for at least one year from the index date. Two primary reviewers adjudicated LC status (M.D. and I.Y.C.). The third and fourth reviewers (J.C.O. and P.C.K.) were consulted to achieve consensus when needed. Demographics, comorbidities, medications, hospitalization, and laboratory results were digitally extracted from the Mayo Clinic Unified Data Platform and our EHR system (Epic Systems, Inc. Verona, WI) using rapidSQL (Idera, Inc., Austin, TX). Manual chart reviews were conducted to confirm SARS-CoV-2 test date and reason for testing, last clinic encounter date, index COVID-19 disease severity, complications, comorbidities (e.g., inflammatory bowel disease, immunosuppression, solid organ transplantation), and 30-day use history of antibiotics, proton pump inhibitors, or immunosuppressive medications. Further information on the antibiotic type, duration, route of administration, and reason for use was obtained. Data were stored and managed using REDCap electronic data capture tools hosted at Mayo Clinic^33^.

We classified COVID-19 vaccination status as fully vaccinated, first dose, and unvaccinated. Patients were considered fully vaccinated two weeks after they completed a two-dose mRNA COVID-19 live attenuated vaccine series (BNT162b2 [Pfizer-BioNTech], mRNA-1273 [Moderna]) or a single-dose viral vector live attenuated vaccine (Ad26.COV2.S [Janssen])^34^. COVID-19 disease severity was adopted from the NIH COVID-19 treatment guidelines and consisted of five categories: (1) asymptomatic or pre-symptomatic, (2) mild illness, (3) moderate illness, (4) severe illness, and (5) critical illness^35^. Comorbidities were classified and standardized using the unweighted Charlson Comorbidity Index (CCI) score using the “comorbidity” package in R (v4.3.3; R Core Team 2024)^36,37^.

### Statistical analysis

#### Descriptive statistics

Descriptive statistics were reported using frequencies and proportions for categorical data and mean, median, standard deviation (SD), and interquartile range (IQR) for continuous variables. Categorical data were analyzed using the chi-square (χ2) test or Fisher’s exact test, while group comparisons for continuous variables were conducted using Student’s *t*-test or Wilcoxon–Mann–Whitney test. Assessment of clinical data completeness revealed that missing data were minimal, constituting less than 0.07% of the entire dataset per respective variables used to adjust microbiome analysis and as model input. Missing values were imputed using mean, median, or mode methods, depending on the data type. Statistical significance was set at *p*<0.05, and all statistical analyses were performed using R.

#### Microbiome data analysis

Microbial alpha and beta diversity analyses were conducted using the R “vegan” (v2.6-6.1) and “GUniFrac” (v1.8) packages after rarefaction^38,39^. Chao1 species richness estimator and Shannon indices were used for alpha diversity analysis while Bray-Curtis (BC) dissimilarities and generalized UniFrac (GUniFrac) distance were used for beta diversity analysis. One-way ANOVA and PERMANOVA were used for testing the group association with alpha and beta diversity, respectively^39^. Statistical significance for diversity analysis was set at *p*<0.05. For post hoc analysis of beta diversity, “RVAideMemoire” (v0.9-83-7)^40^ was employed for pairwise comparisons with false discovery rate (FDR) correction. To investigate the relationship between microbial diversity and LC group membership, redundancy analysis (RDA), a form of constrained ordination, was conducted using the “vegan” package^38^. This analysis involved assessing a matrix of interval-scaled data (species abundance) against a matrix of covariates through singular value decomposition. Procrustes analysis was conducted on longitudinal microbiome data to inspect any changes in microbial beta diversity between T1 (acute phase) and T2 (post-acute phase) samples^41^. To quantify and compare the magnitude of microbiome changes over time across three cohorts, we extracted pairwise Procrustes by sample and calculated Euclidean distances. Differential taxa abundance analysis was performed using a linear model-based permutation procedure in the “ZicoSeq” function of the “GUniFrac” package^39,42^. The FDR-adjusted *p*-value threshold was set at 0.1 (i.e., allowing on average 10% false discoveries in the results to identify taxa with moderate effects). Approaches for data pre-processing and handling are described in **Supplemental Methods**.

### Software and algorithms

#### Applying machine learning models to predict LC

Machine learning (ML) algorithms — logistic regression with L1 regularization, and artificial neural network (ANN) — were employed to model LC in our study population. The goal of the models was to classify LC (n=80) from the COVID-19-positive cohort (n=380). Input data types included species-level (n=5671) and genus-level (n=1456) relative abundance matrices, and clinical variables, which included age, sex, body mass index (BMI), baseline comorbidity score (CCI), index COVID-19 severity, SARS-CoV-2 vaccination status, immunosuppressive medication use, and 30-day antibiotic use. Class weights were calculated using “scikit-learn” (v1.2.2) based on the frequencies of labels (0, 1), with weights assigned inversely proportional to prevent bias toward overrepresented classes^43^. Model development and validation were performed using nested cross-validation with architecture of an inner loop for hyperparameter tuning using “GridSearchCV” function^43^ and an outer loop for validating each model’s performance. This setup was repeated 10 times on random seeds for reproducibility. Five-fold cross-validation was applied to the inner and outer loop and performance metrics, including accuracy and the weighted F1 score, which were calculated for each model configuration to select the optimal parameters. Feature importance was quantified using SHapley Additive exPlanations (SHAP)^44^ values and log-odds, calculated to assess the impact of each feature on model predictions. All models were developed using Python (Python Software Foundation, Beaverton, OR). Details in model development, cross-validation, and interpretation are described in **Supplemental Methods**

#### Modeling LC disease heterogeneity

To identify potential disease subphenotypes in LC, 18 symptoms observed in more than 10% of LC patients were extracted. Multiple Correspondence Analysis (MCA) was conducted to reduce its dimensionality and derive coordinates using “factoextra” (v1.0.7)^45^, and the resulting MCA factor plot (**Fig. S1A**) was examined to assess data structure and discernable clustering tendencies. A heatmap⁴⁶ with hierarchical and *k*-means clustering (n=999) was then generated on symptom matrix, using a binary dissimilarity metric to further delineate symptom clusters.

To determine the optimal number of clusters (*k*), binary distance matrix was computed, followed by classical multidimensional scaling (MDS) to reduce the noise in data observed in MCA plots. The first three MDS dimensions, accounting for 47.17% of total variance, were used to calculate the within-cluster sum of squares (WCSS) for *k* from 2 to 10. An elbow was observed at *k*=4 (**Fig. S1B**), balancing cluster coherence and parsimony. To further support the findings with elbow technique, mean silhouette widths s(*i*), were calculated for iterated *k* values (2-10) using “cluster” (v2.1.6)^47^ package which showed a peak at *k*=4 (**Fig. S1C**).

## Data availability statement

Data tables, and analytic methods will be published and available to other researchers in supplemental materials. The microbiome data is publicly available on NCBI’s BioProject database with BioProject ID PRJNA1167328. The SARS-CoV-2 genome sequences from this study have been deposited in GISAID’s EpiCoV database under GISAID identifier EPI_SET_240908tm.

## Acknowledgments

We are grateful to all the study participants for providing samples and their involvement in the study. We also thank Rochelle M. Fabian, Wilkes Quantina, Ilyse Nelson, and Savannah Van Eycke for their roles as study coordinators leading patient enrollment. Lastly, we appreciate the contributions of the Center for Individualized Medicine operations managers in ensuring the successful completion of this study. This project was supported by funding from the Center for Individualized Medicine (CIM), a division in Mayo Clinic. APA received research funding from NSF-IIS-2041339. GKG received funding from NIH NIGMS R35GM149270. PCK is supported by NIH NIDDK R01DK114007.

## Author Contributions

P.C.K. and K.N.L. conceived the project. I.Y.C., L.Y. J.C., and S.J. contributed to the formal data analysis and visualization. I.Y.C. wrote the original draft. I.Y.C., R.T.M., M.D. contributed to data curation. P.C.K., I.Y.C., R.T.M., M.D., L.R.H., A.K.K., J.P.S., and E.L. conducted investigation. P.C.K., J.C.O., C.A., A.P.A. and G.K.G. supervised the project. P.C.K., T.M.V.G., J.J.H., administered the clinical investigation. All authors reviewed, and approved the final manuscript.

## Competing interests

Authors declare no competing interest.

## Materials & Correspondence

Correspondence to Purna C. Kashyap

## Supplementary Figures

**Supplemental Figure 1.**
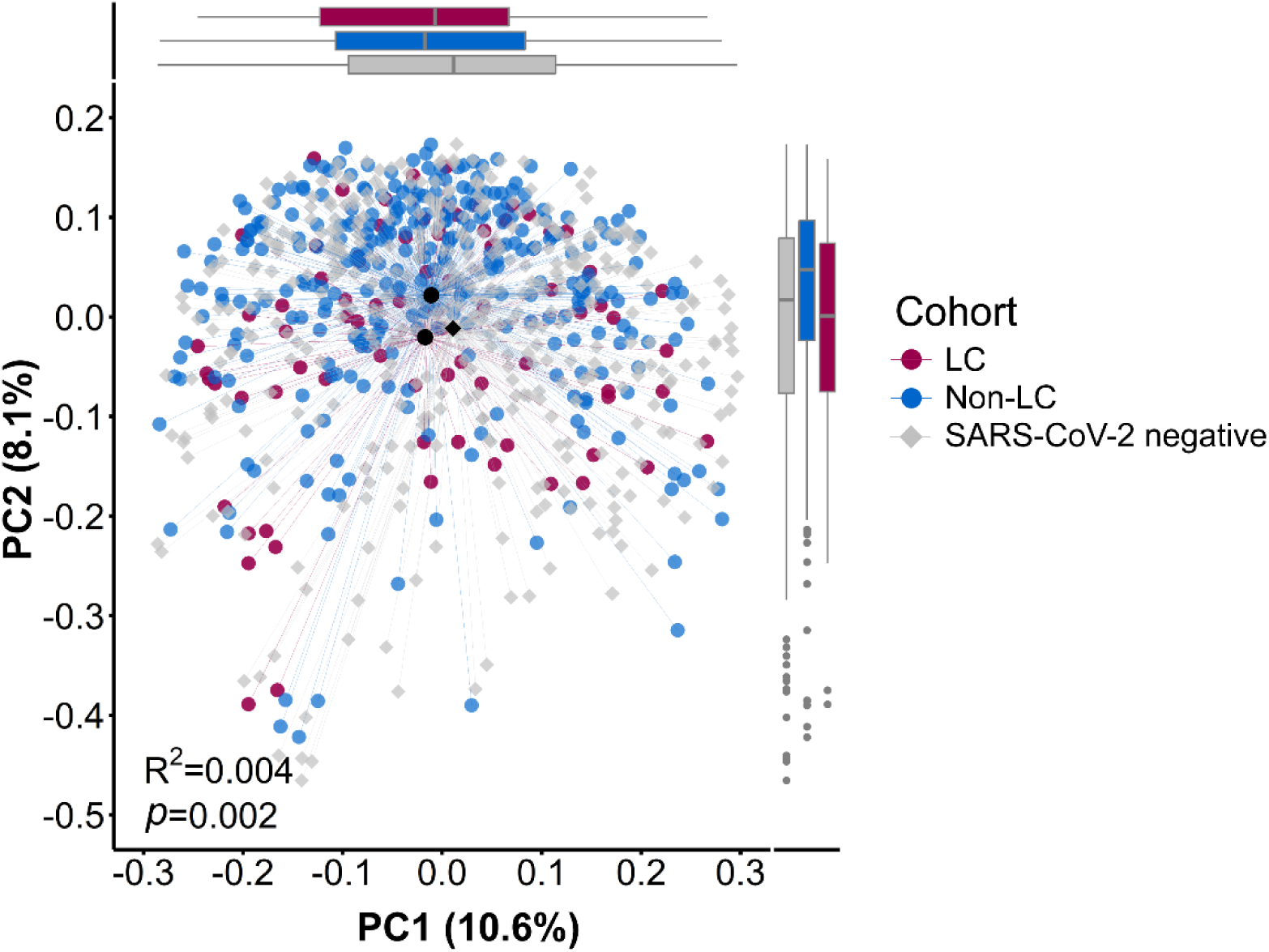
The gut microbiome is altered during acute infection in patients with LC. Similar to Bray–Curtis distances, generalized UniFrac distances at the species level showed significantly different microbiome composition during acute infection across LC, non-LC, and SARS-CoV-2-negative groups (PERMANOVA, n=999, R^2^=0.004, *p*=0.002) after adjusting for age, sex, baseline comorbidity score, and 30-day antibiotic use. Post hoc analysis found that the microbiome composition of the LC group differed significantly from the non-LC group (pairwise comparisons using PERMANOVAs, n=999, FDR-adjusted *p*=0.006) and SARS-CoV-2-negative group (FDR-adjusted *p*=0.038).

**Supplementary Figure 2.**
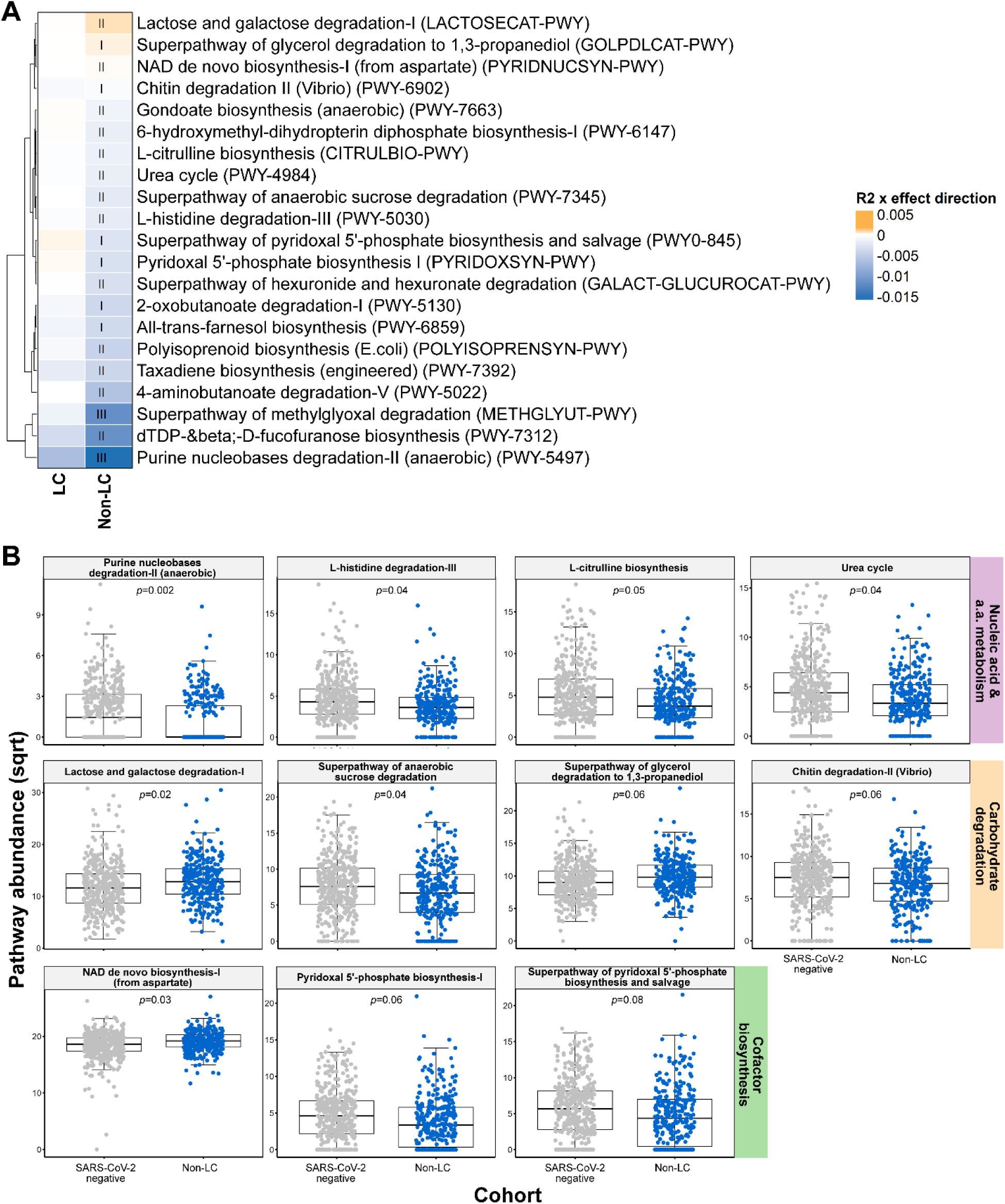
Comparative analysis of microbial functional pathway abundances. **(A)** Differential abundance analysis of microbial pathways using ZicoSeq highlighting nominal associations between LC (n=80) and non-LC (n=299) cohorts compared to SARS-CoV-2-negative controls (n=419) adjusted for clinical confounders. Level of significance are represented by Roman numerals: I, FDR-adjusted *p*<0.1; II, *p*<0.05; and III, for *p*<0.01. **(B)** Comparison of individual microbial pathways. In all panels, boxplots represent data from non-LC and SARS-CoV-2-negative cohorts. The central line indicates the median, and the box represents the interquartile range (IQR). Whiskers extend to data points within 1.5 × IQR. FDR-adjusted *p*-values were generated using ZicoSeq. Pathway types were determined using ontology-based MetaCyc^48^ pathway classification.

**Supplementary Figure 3.**
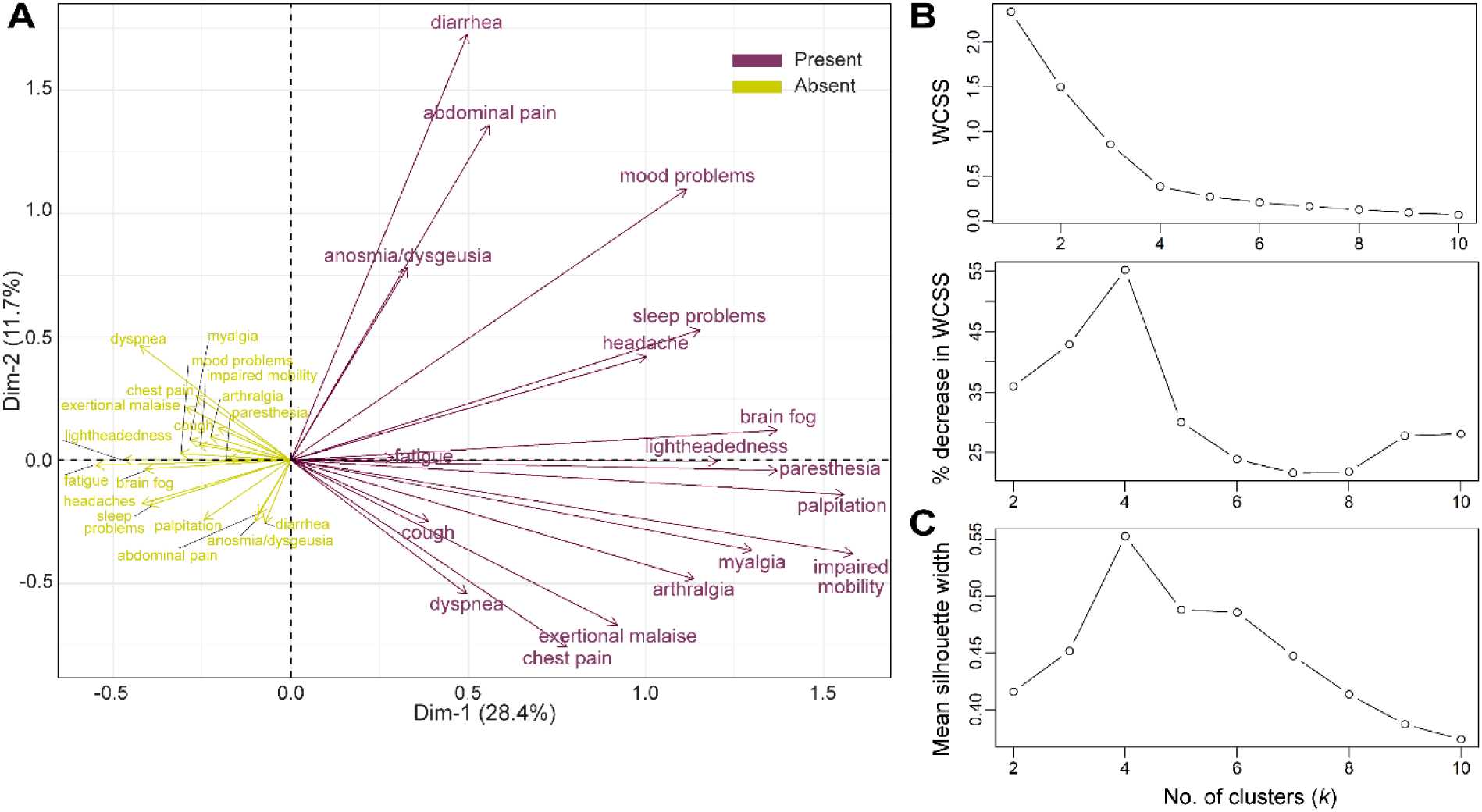
Determining the optimal number of clusters for analyzing disease heterogeneity. **(A)** Multiple correspondence analysis (MCA) plot of variables (symptoms) illustrating the relationships between symptoms categorized as “Present” (purple) and “Absent” (yellow) in the binary dataset. Symptoms clustering in similar regions indicate shared patterns or associations. The length and direction of each arrow indicate the strength and orientation of the symptom’s contribution to the variability captured in first two dimensions. **(B)** The within-cluster sum of squares (WCSS) for *k*-means clustering of LC symptoms was plotted against numbers of clusters (upper panel), along with the corresponding percentage decrease in WCSS (lower panel). An “elbow” in the plot was seen at *k*=4, suggesting that four clusters optimally balance cluster cohesion and parsimony. **(C)** Mean silhouette widths s(*i*) were calculated for *k* values ranging from 2 to 10. The silhouette width for each symptom measures how well it fits within its own cluster compared to others; with higher values closer to one indicating the no. of cluster providing the more cohesive grouping of the data. The average silhouette width peaks at *k*=4.

## Supplementary Tables

**Supplemental table 1.** Clinical characteristics of the study population.

|  |  |  |  |  |
| --- | --- | --- | --- | --- |
|  | LC (n=80) | Non-LC (n=300) | SARS-CoV-2-negative (n=419)<sup>1</sup> | *P* |
| Chronic pulmonary disease | 21 (26) | 32 (11) | 68 (16) | 0.004 |
| Diabetes mellitus | 9 (11) | 22 (7) | 37 (9) | 0.5 |
| Chronic kidney disease | 3 (4) | 24 (8) | 30 (7) | 0.4 |
| Moderate to severe liver disease | 0 | 4 (1) | 0 | - |
| Myocardial infarction | 2 (2.5) | 3 (1) | 9 (2) | 0.4 |
| Rheumatological disease | 4 (5) | 7 (2) | 20 (5) | 0.2 |
| Metastatic cancer | 3 (4) | 7 (2) | 13 (3) | 0.6 |
| Inflammatory bowel disease | 4 (5) | 8 (3) | 10 (2) | 0.4 |
| Transplant history | 2 (2.5) | 6 (2) | 4 (1) | 0.3 |
| Solid organ transplant | 2 (2) | 3 (1) | 3 (1) |  |
| Bone marrow transplant | 0 | 3 (1) | 1 (0.2) |  |
| Proton pump inhibitor use | 20 (25) | 58 (19) | 95 (23) | 0.4 |
| Immunosuppressive medication<sup>2</sup> | 7 (9) | 13 (4) | 32 (8) | 0.2 |
| Glucocorticoids | 1 (1) | 3 (1) | 10 (2) |  |
| Antimetabolites | 1 (1) | 1 (0.3) | 8 (2) |  |
| Cancer chemotherapeutics | 1 (1) | 1 (0.3) | 6 (1) |  |
| Calcineurin inhibitors | 2 (2.5) | 2 (1) | 1 (0.2) |  |
| Antiproliferative agents | 1 (1) | 5 (2) | 4 (1) |  |
| Biologics | 3 (4) | 6 (2) | 5 (1) |  |
| Other | 1 (1) | 1 (0.3) | 2 (0.5) |  |

Data is represented as n (%).
*P*-values were generated using Pearson's Chi-squared test, or Fisher's exact test.
<sup>1</sup>Comorbidities and medication use history of three SARS-CoV-2-negative controls are missing (same patients across variables).
<sup>2</sup>Immunosuppressive medication use within 30 days prior to SARS-CoV-2 test date. Glucocorticoid use was defined as the equivalent to >7.5 mg total daily prednisone for a minimum of four weeks.

**Supplemental table 2.** Distribution of symptom types at the time of SARS-CoV-2 testing in the symptomatic cohort.

|  |  |  |  |
| --- | --- | --- | --- |
|  | LC (n=79) | Non-LC (n=268) | SARS-CoV-2-negative (n=234) |
| Symptom type, index testing, n (%) |  |  |  |
| Cough | 57 (72) | 168 (63) | 72 (31) |
| Headache | 46 (58) | 146 (54) | 73 (31) |
| Myalgia | 36 (46) | 117 (44) | 44 (19) |
| Rhinorrhea/nasal congestion | 38 (48) | 107 (40) | 20 (8.5) |
| Fever | 26 (33) | 92 (34) | 41 (18) |
| Fatigue | 35 (44) | 72 (27) | 42 (18) |
| Chills | 24 (30) | 81 (30) | 36 (15) |
| Sore throat | 23 (29) | 77 (29) | 75 (32) |
| Dysgeusia | 26 (33) | 61 (23) | 6 (3) |
| Anosmia | 21 (27) | 61 (23) | 3 (1) |
| Dyspnea | 26 (33) | 46 (17) | 32 (14) |
| Chest distress | 29 (37) | 36 (13) | 16 (7) |
| Diarrhea | 20 (25) | 44 (16) | 38 (16) |
| Nausea | 17 (22) | 24 (9) | 45 (19) |
| Sputum production | 15 (19) | 25 (9) | 14 (6) |
| Lightheadedness | 16 (20) | 22 (8) | 7 (3) |
| Abdominal pain | 8 (10) | 12 (4) | 11 (5) |
| Vomiting | 5 (6) | 7 (3) | 19 (8) |
| Missing | 11 (14) | 29 (11) | 67 (29) |

**Supplemental table 3.** Additional stool characteristics and SARS-CoV-2 genotype of the study population, all cohorts.

|  | Timepoint-1 |  |  | Timepoint-2 |  |  |
| --- | --- | --- | --- | --- | --- | --- |
|  | LC<br>(n=80) | Non-LC<br>(n=300) | SARS-CoV-2-negative<br>(n=419) | LC<br>(n=27) | Non-LC<br>(n=73) | SARS-CoV-2-negative<br>(n=48) |
| Bristol score |  |  |  |  |  |  |
| 1 | 4 (5) | 5 (2) | 12 (3) | 2 (7) | 2 (3) | 1 (2) |
| 2 | 7 (9) | 15 (5) | 29 (7) | 1 (4) | 6 (8) | 3 (6) |
| 3 | 9 (11) | 38 (13) | 64 (15) | 5 (18.5) | 5 (7) | 8 (17) |
| 4 | 23 (29) | 94 (31) | 124 (30) | 7 (26) | 23 (31.5) | 12 (25) |
| 5 | 4 (5) | 16 (5) | 23 (5.5) | 2 (7) | 5 (7) | 2 (4) |
| 6 | 7 (9) | 31 (10) | 38 (9) | 0 | 8 (11) | 5 (10) |
| 7 | 3 (4) | 6 (2) | 2 (0.5) | 0 | 0 | 0 |
| Missing | 23 (29) | 95 (32) | 127 (30) | 10 (37) | 24 (33) | 17 (35) |
| SARS-CoV-2 genotype |  |  |  |  |  |  |
| AY.100 | - | 2 (1) | - | - | - | - |
| AY.103 | - | 3 (1) | - | - | - | - |
| AY.25 | - | 1 (0.3) | - | - | - | - |
| AY.3 | - | 2 (1) | - | - | - | - |
| AY.44 | - | 1 (0.3) | - | - | - | - |
| AY.47 | 1 (1) | - | - | - | - | - |
| AY.54 | - | 1 (0.3) | - | - | - | - |
| B.1 | - | 1 (0.3) | - | - | - | - |
| B.1.1.222 | - | 1 (0.3) | - | - | - | - |
| B.1.2 | 2 (2.5) | 9 (3) | - | - | - | - |
| B.1.234 | 1 (1) | 1 (0.3) | - | - | - | - |
| B.1.239 | - | 1 (0.3) | - | - | - | - |
| B.1.243 | - | 1 (0.3) | - | - | - | - |
| B.1.311 | 1 (1) | - | - | - | - | - |
| B.1.427 | - | 1 (0.3) | - | - | - | - |
| B.1.429 | - | 1 (0.3) | 1 (0.2) | - | - | - |
| B.1.565 | - | 1 (0.3) | - | - | - | - |
| B.1.617.2 | - | 5 (2) | - | - | - | - |
| Unassigned (e.g., poor coverage, high ambiguity, conflicting lineage) | 1 (1) | 3 (1) | 2 (0.4) | - | 1 (1) | - |
| Categorical variables are reported as n (%).<br>Genomic lineages of SARS-CoV-2 sequences were determined using the Phylogenetic Assignment of Named Global Outbreak Lineage (Pangolin) |  |  |  |  |  |  |

**Supplementary table 4.** Overview of studies in gut microbiome and long COVID with metagenomics stool sampling.

|  |  |  |  |  |
| --- | --- | --- | --- | --- |
| Year, author | SARS-CoV-2 positive cohort size (no. of patients) | Hospitalization status | Sequencing method | Timing of stool collection |
| Liu Q et al (2022)<sup>1</sup> | 133 | Inpatient during acute infection | Shotgun metagenomic sequencing | At hospital admission, 3 months<br>6 months after respiratory SARS-CoV-2 viral clearance. |
| Liu Q, Mak JWY, Su Q, et al. (2022)<sup>2</sup> | 106 | Inpatient during acute infection | Shotgun metagenomic sequencing | At hospital admission, 1 month and 6 months after the hospital discharge |
| Zhang et al. (2023)<sup>3</sup> | 187 | Inpatient during acute infection | 16S rRNA gene sequencing (V3–V4) | 12 (+/- 1) months after the hospital discharge |
| Su Q et al. (2024)<sup>4</sup> | 196 | Inpatient during acute infection | Shotgun metagenomic sequencing | During hospitalization |

## Supplementary Methods

### Microbial DNA extraction

Briefly, PowerBead Pro plates were loaded in a TissueLyser II for homogenization before loading samples onto a QIAcube HT for automated DNA extraction. This protocol was selected because it includes both chemical and mechanical lysis, which have been shown to be important for accurate recovery of abundances of difficult-to-lyse bacteria (e.g., some Gram-positive species), and allows for automation, which has been shown to reduce batch variability effects. DNA was quantified using Qubit.

### Statistical analysis

#### Microbiome data analysis

For microbiome data pre-processing, 10% prevalence and 0.2% maximum abundance filters were applied. Outliers were handled using winsorization, capping the top 3% of data points, thereby minimizing the influence of extreme values. Zeroes were imputed through posterior sampling, which was conducted 25 times to reduce the variability of results. Data were transformed using multiple link functions including fourth root, square root, and three-quarters power transformations (unless otherwise specified) to model non-linear relationships between clinical covariates and microbial counts. The statistical significance was assessed using permutation-based false discovery rate (FDR). The analysis also included reference-based multiple-stage normalization to address compositional effects, using six normalization stages with a reference set comprising 50% of the samples. Differential abundance analysis was conducted using ZicoSeq package in R [1]. Metagenomics functional pathways were analyzed using the same pre-processing steps and tools. Data visualization for microbial taxa and functional pathway comparisons were adopted from standard biostatistical tools [2-4].

### Software and algorithms

#### Data pre-processing

One-hot encoding was applied to transform categorical variables from the clinical data into a binary matrix. Numerical data from both datasets were normalized by scaling the values between 0 and 1 using MinMaxScaler. To normalize microbiome abundance matrices, we applied scaling between 0 and 1. No feature selection was applied to clinical variables, and no pre-filtering was applied to microbiome data.

#### Model development and nested cross-validation

##### Logistic regression with L1 regularization

Logistic regression from scikit-learn (v1.2.2) was employed with the ‘liblinear’ solver [5]. The model was configured to incorporate the computed class weights to correct for unbalanced classes. Hyperparameter tuning was performed via GridSearchCV during nested cross-validation, optimizing the regularization strength parameter *C* over 100 evenly spaced values on a logarithmic space, covering multiple orders of magnitude from 10^-3^ to 10^1^ (0.001 to 10). The scoring metrics for model optimization were accuracy and weighed F1 score.

##### Artificial neural network

Models were constructed using TensorFlow’s Keras API (v2.14.0) [6]. The model architecture included three dense layers, configured with 64, 32, and 1 neuron(s), and two dropout layers. ReLU activation function was employed in the first two layers and a sigmoid function in the output layer for binary classification. Models were trained using the Adam optimizer with binary cross-entropy loss, over 50 epochs with a callback function from Keras library to stop training after three epochs with no reduction in loss. The optimization process was conducted via a structured grid search within a nested cross-validation framework, specifically targeting the dropout rates to control for feature selection. accuracy or the weighted F1 scores. A grid search was performed to optimize accuracy and weighted F1 score. Dropout rates (fraction of input units to drop) were tested at intervals of 0.2 (0, 0.2, 0.4, 0.6) to reduce computational demand compared to using smaller increments such as 0.1.

#### Model interpretation

To determine the impact of individual bacterial taxa on model output from logistic regression, coefficients were derived from log-odds. To interpret ANN predictions and identify genera with the greatest impact on the model’s classifications, Shapley values were calculated using “GradientExplainer” function from SHAP (SHapley Additive exPlanations, v0.43.0) package [7]. For these experiments, we chose the models integrating species-level microbiome data and clinical variables, given their slightly better performance compared to other models. To assess consistency and repeatability of the top 20 most influential predictors, the analysis was structured around 10 random seeds coupled with 5-fold cross-validation in the outer loop, which resulted in 50 runs per model. Statistically, each predictor was treated as having an equal probability of being selected as one of the top 20 most influential features The discrete random variable X=1 indicated a feature’s selection as one of the top 20 most influential predictors vs. X=0 otherwise. Given the large number of predictors (n=5,675) and low probability of any single predictor being selected, Poisson distribution was employed to approximate the binomial distribution. Z-scores were calculated for each predictor to assess deviation from the expected frequency, aiding in their ranking based on occurrence. A heatmap was generated to visually display the top features across the three models.

*n=*50 (Bernoulli trials)

*x=*no. of appearance in top 20 for each predictor

*p=*probability of the event (20/5675)

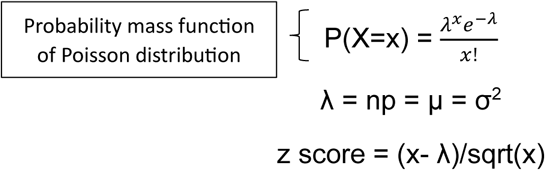

## Notes

### Competing Interest Statement

The authors have declared no competing interest.

